# Clathrin-coated Pits are Shredded Organelles Presorting Receptors in the Plasma Membrane

**DOI:** 10.1101/491670

**Authors:** Do-Hyeon Kim, Yonghoon Kwon, Hyeong Jeon An, Kai Zhou, Min Gyu Jeong, Soyeon Park, Yeonho Chang, Nam Ki Lee, Sung Ho Ryu

## Abstract

A myriad of receptor types crowds the plasma membrane of cells. Here, we revealed that clathrin-coated pits (CCPs) presort receptors to modulate receptor activation on the plasma membrane. We visualized individual molecules of receptors, including epidermal growth factor receptor (EGFR), ErbB2, transferrin receptor (TfR), and beta2-adrenergic receptors (β2-AR), inside a single CCP in a living cell using single-molecule diffusivity-based colocalization analysis. The spatially distinct subsets of CCPs selectively allocated for these receptors were observed in a resting state. The EGFR pre-allocated CCP subset was partially shared with that for ErbB2, whereas the EGFR and TfR pre-allocated CCP subsets were mutually exclusive. PICALM was necessary for the pre-allocation of CCPs for EGFR. Furthermore, EGFR dimerization was markedly elevated inside the pre-allocated CCP subset after dynamin recruitment. The pre-sorting function of CCPs provides an efficient mechanism to control competition and cooperation between receptors on the crowded plasma membrane.

**One Sentence Summary:** Clathrin-coated pits are small-sized organelles pre-sorting receptor kinds to modulate receptor activation on the crowded plasma membrane.

## Introduction

Receptors sense extracellular cues and send appropriate signals inside the cell (*1*). A myriad of receptors co-exist in the crowded plasma membrane to cope with a capricious environment around the cell (*2*). In general, these receptors are activated upon binding of their own ligands, which is accompanied by receptor dimerization/oligomerization (*3, 4*). However, the precise nature of receptor activation in the membrane of living cells is not fully understood (*5, 6*). There have been numerous reports indicating the existence of signaling platforms or hotspots, such as lipid rafts, non-raft microdomains, and membrane skeletons, which may control receptor activation modes and differential signaling pathways (7–10).

Clathrin-mediated endocytosis (CME) is the one of the major internalization pathways for various receptors, including enzyme-linked receptors, G protein-coupled receptors and ion channels (*11, 12*). Upon ligand-induced activation, these receptors undergo internalization via a dynamic endocytic machinery termed clathrin-coated pits (CCPs) (*13*). Rapid internalization of the activated receptors diminishes their abundance on the plasma membrane, which results in desensitization to the ligand and attenuation of receptor signaling (*14*). Recently, it has been shown that the residence time of receptors inside CCPs controls their signaling pathways (*15*) and receptors remain in an active state in endosomes after CME producing endosomal signaling (*16*). These findings indicate that CCPs actively participate in receptor signaling, raising the question of whether CCPs have a distinct function from their endocytic role as modulators of receptor signaling.

Here, we found that the distinct subsets of CCPs are pre-allocated for different kinds of receptors, including epidermal growth factor receptor (EGFR), ErbB2, transferrin receptor (TfR), and beta2-adrenergic receptor (β2-AR), on the crowded plasma membrane in a resting state by directly visualizing individual receptor molecules inside CCPs in living cells using diffusion-based molecular colocalization analysis. Furthermore, we revealed that the pre-allocated CCP subset for EGFR coordinates its activation by facilitating EGFR dimerization, which demonstrates that CCPs serve a sorting platform to modulate receptor signaling.

## Results

### Diffusion-based molecular colocalization between EGFR and CCSs in living cells

EGFR signaling dependent on CME was previously reported (*17*). In search of the onset of EGFR signaling triggered by CME, we investigated whether EGFR in a resting state (serum starvation) interacts with clathrin-coated structures (CCSs) on the plasma membrane. The colocalization between CCS and EGFR after EGF-induced activation on the plasma membrane has been analyzed using total internal reflection fluorescence microscopy (TIRFM) because activated EGFR is typically clustered inside CCSs, which produces a high contrast of fluorescence intensity inside CCSs over the plasma membrane. However, in the resting state, EGFR is well-distributed across the plasma membrane at a high density, which interferes with colocalization analysis between individual EGFR molecules and CCSs even at the 20-nm resolution of super-resolution microscopy (Figure 1A). We resolved this problem by adding an extra dimension, diffusivity, to the space and time dimensions, based on the concept that the diffusivity of two molecules must be the same if they physically interact with each other (*18–20*), which provides a criterion for discriminating true from falsely colocalized molecules.

**Figure 1.**
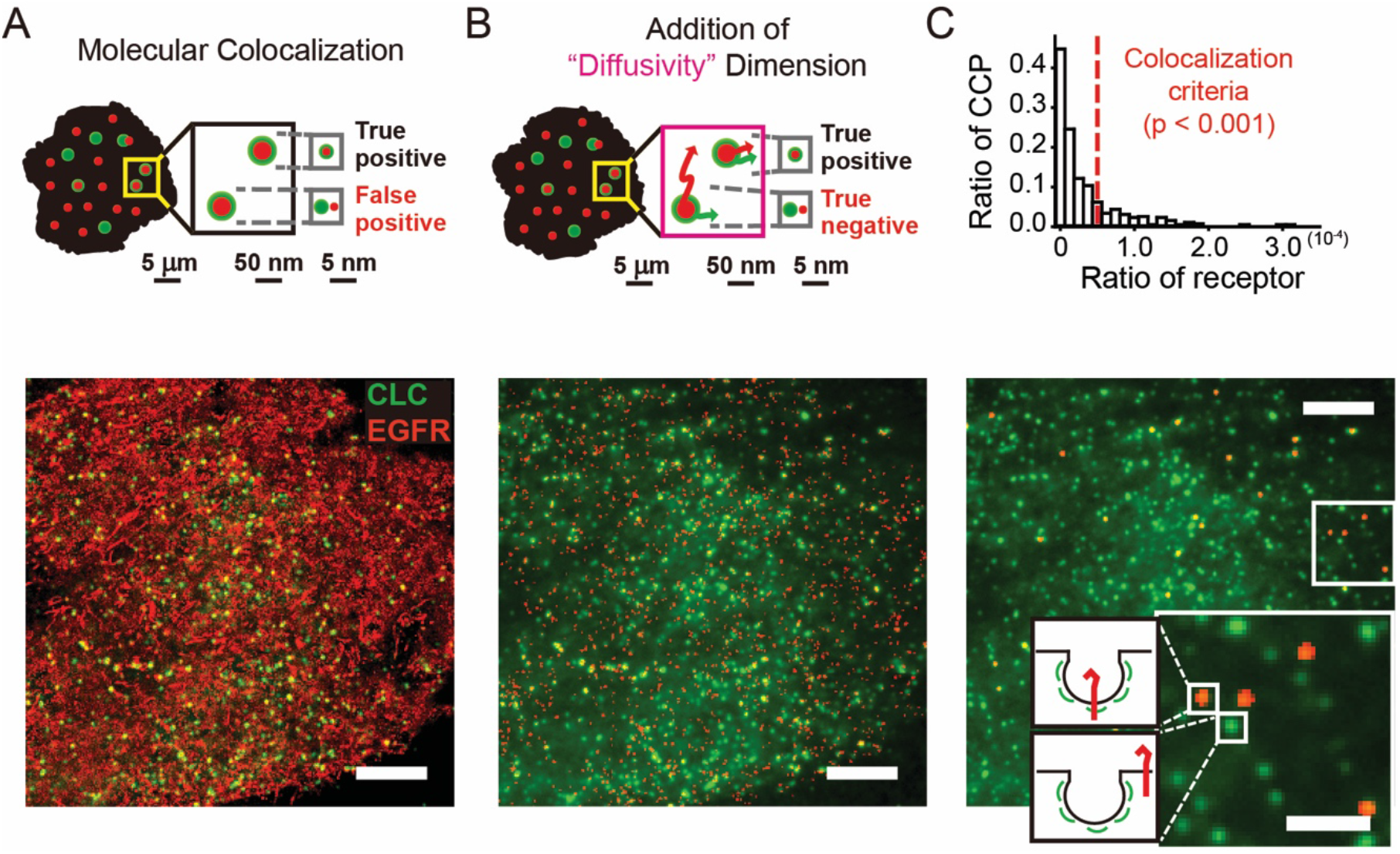
Diffusion-based molecular colocalization between EGFR and CCP. (A) A schematic concept of molecular colocalization based on super-resolution microscopy (*upper panel*). An overlay between the PALM image of EGFR-mEos3.2 (red) and the TIRF image of CLC-EGFP (green) (*lower_panel)*. (B) A schematic concept of diffusion-based molecular colocalization (*upper panel*). An overlay between the PALM image of EGFR-mEos3.2 matching the diffusivity of CCP (red) and the TIRF image of CLC-EGFP (green) (*lower panel*). (C) A histogram for the individual CCPs with respect to the number of spatiotemporally colocalized EGFR matching the diffusivity of the CCPs. The colocalization criteria (p < 0.001) is shown as a red dashed line (*upper panel*). Diffusion-based molecular colocalization image for EGFR and CCP. EGFRs specifically contained in CCPs were displayed (red) in the TIRF image of CLC-EGFP (green). The inset exhibits the magnified image of the selected region (a white box) (*lower panel*). Scale bars, 5 μm (A, B and C) and 1 μm (the inset of C).

To implement the diffusion-based molecular colocalization, we tracked singlemolecule endogenous EGFR labeled by the Fab fragment of anti-EGFR antibody conjugated with Alexa Fluor 647 in BSC1 cells that stably express EGFP-tagged clathrin light chain (CLC) using TIRFM (Figure S1A-C). Individual CCSs were tracked simultaneously with EGFR in the same cell with two different EM-CCDs at a frame rate of 50 Hz for EGFR and of 1 Hz for CCSs corresponding to their respective intrinsic diffusivities (Figure S1D-S1I and Movie S1). There was no sign of photodamage to the cells after the imaging (Figure S1J and S1K). The short-time diffusion coefficients of EGFR and CCS trajectories were determined by mean squared displacement (MSD_EGFR_ = 4Dt, 0 < t < 200 ms, MSD_CLC_ = 4Dt, 0 < t < 5s) with time lags of 20 ms and 1 s, respectively. The images obtained with two EM-CCDs were registered using nanoparticles yielding ~9.4 nm resolution, which exerted a marginal effect on colocalization between EGFRs and CCSs because the localization error of image registration was much lower than that of Alexa Fluor 647 and EGFP (Figure S2A-S2E). We selected the EGFR trajectories exhibiting the same diffusivity to CCSs, which seemed nearly immobilized because CCSs barely moved during the time required for the measurement of EGFR diffusivity (Figure 1B). On average, ~15.6% of total EGFR trajectories were selected, and these trajectories tended to appear in the same positions repeatedly, probably due to small-size clustering, which substantially reduced the spatiotemporal complexity of colocalization between EGFRs and CCSs. Among the EGFR trajectories matching the diffusivity of CCSs, we counted the number of the trajectories colocalizing with CCSs in space and time. Then, we identified the CCSs containing EGFR molecules with a false positive rate of 0.1%, determined by counting the number of CCSs containing EGFRs, whose positions were randomized (Figure S2F and S2G).

The diffusion-based molecular colocalization analysis between EGFR and CCS in a live BSC1 cell showed that an ~13.5% subset of CCSs contained EGFRs in the resting state (Figure 1C), which implies an early involvement of CCSs in EGFR signaling prior to ligand-induced EGFR activation.

### Validation of the diffusion-based molecular colocalization between EGFR and CCSs

Although we statistically verified the specificity of the diffusion-based molecular colocalization between EGFRs and CCSs by calculating a false-positive rate using randomization, further experimental validation of the method was required. Thus, we first used a plasma membrane-targeting peptide (PMT) tagged to mEos3.2, a monomeric photoactivatable fluorescent protein, which enforces mEos3.2 to be localized on the plasma membrane. We observed only ~1.2% of CCSs containing PMT, although PMT-mEos3.2 exhibited an ~34.4% immobilized subpopulation (Figure 2A and 2B). We also tested an AP2 binding-deficient EGFR mutant (EGFR ΔAP2) of which CME is hindered because AP2 is a critical factor for CME and involved from the very early stage of CCS development (*21*). EGFR ΔAP2 exhibited a significantly decreased fraction of CCSs containing EGFR ΔAP2 (~2.9%) compared to wild-type (WT) EGFR (Figure 2C and 2D).

**Figure 2.**
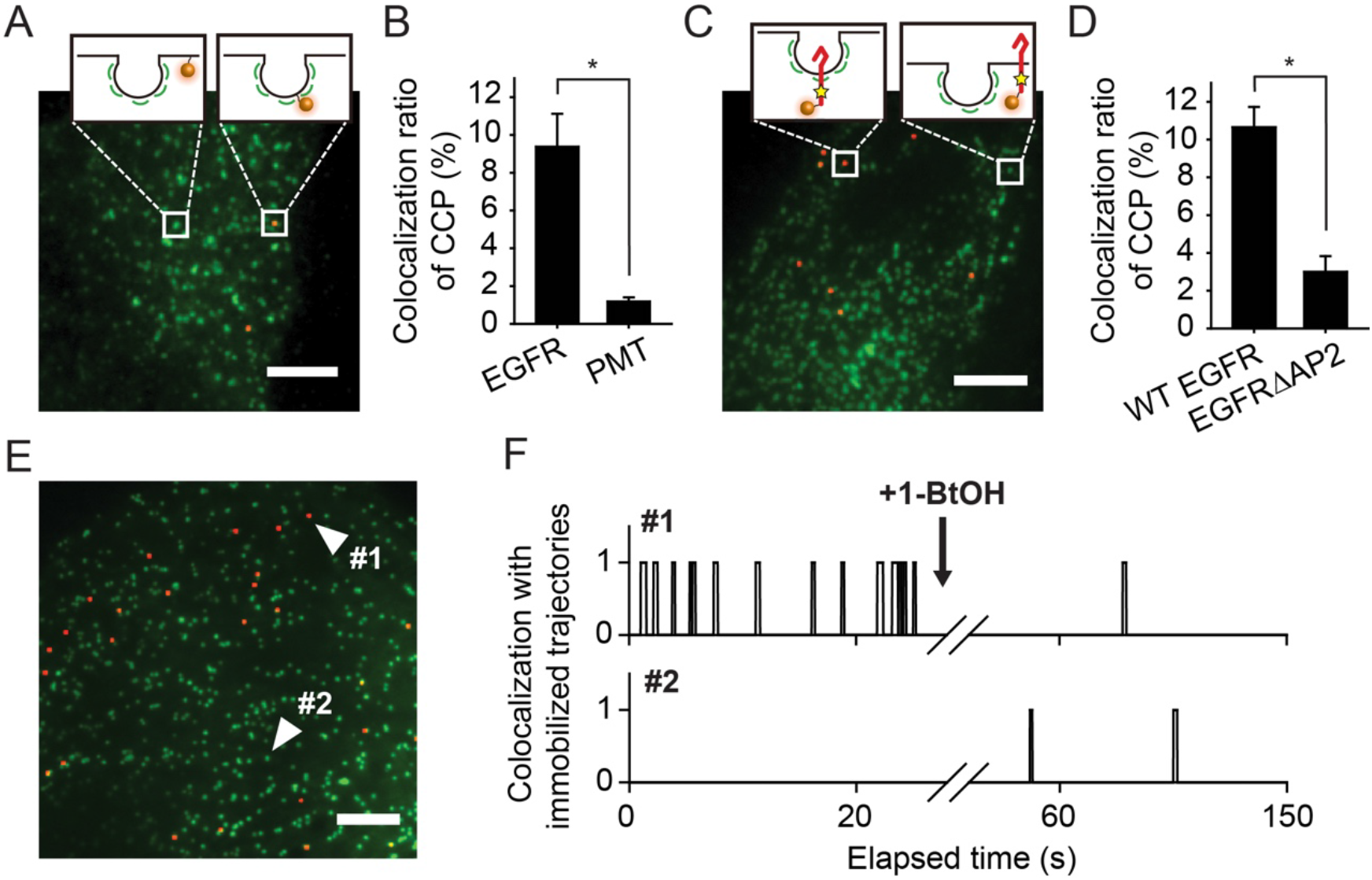
Validation of specificity of the diffusion-based molecular colocalization of EGFR with CCP. Diffusion-based molecular colocalization analysis of CCPs with PMT (A), EGFR ΔAP2 (C), and WT EGFR (E). Histograms represent the colocalization ratio of CCPs with PMT (B) or EGFR ΔAP2 (D). (F) The time profile of colocalization events of the indicated individual CCPs with single-molecule WT EGFR trajectories matching the diffusivity of CCPs before and after the treatment with 1-butanol. Error bars denote s.e.m. at the single-cell level (n > 5). Scale bars, 5 μm. *p < 0.01 (Student’s *t*-test)

Next, we examined the effect of perturbation in CCSs on the colocalized EGFR trajectories. We acutely removed CCSs using 1-BtOH immediately after we acquired the diffusion-based single-molecule colocalization image, and then we analyzed the changes in the colocalized EGFR trajectories (Figure S3) (*22*). After the removal of CCSs, the multiple reappearances of immobilized EGFR trajectories were no longer detected at the positions where CCSs has existed before 1-BtOH treatment (Figure 2E and 2F). Altogether, these results indicated that the CCS subset containing EGFR in the resting state was specifically visualized.

### CCPs Are Pre-allocated For Different Kinds of Receptors on a Plasma Membrane

We next investigated whether the existence of the CCS subset containing EGFR in the resting state originates from the temporally heterogeneous lifecycle of CCSs or the spatially distinctive subset formation of CCSs in living cells. Pitstop 2 is a cell-permeable inhibitor targeting a clathrin terminal domain, which stalls the lifecycle of CCSs by blocking their maturation (*22*). Upon incubation with Pitstop 2, the lifecycle of almost every CCS in a single cell was stalled as a pit form and the lifetime of these CCPs lasted longer than 5 min (Figure S4). In the presence of Pitstop 2, the CCP subset containing EGFR was still observed at ~12.2 % of the total CCPs (Figure 3A and 3B). Interestingly, the status of EGFR containment for individual CCPs was maintained over time: the CCP subset with EGFR tended to keep EGFR over time, whereas the CCP subset without EGFR tended to keep no EGFR. The conditional probability of the CCP subset for EGFR at 0 min overlapping the subset after 5 min was 0.77, and the independence test confirmed their strong dependency (~6.25) (Figure 3B). This result might be derived from a pitfall that EGFR could not dynamically access other CCPs at the plasma membrane if the pits were closed by Pitstop 2. However, it was previously reported that these pits remain open after Pitstop2 treatment (*22*). We further corroborated the dynamic accessibility of single-molecule EGFR to CCPs by analyzing the long trajectories of EGFR to spot a transient CCP-trapping process (*23*). We observed that EGFR enters a CCP, resides in it for an average of ~1.1 s, and then exits (Figures 3C and Movie S2). The existence of the spatially distinctive CCP subset selectively allocated for EGFR indicates that the fates of maturating CCPs are predetermined in terms of EGFR containment.

**Figure 3.**
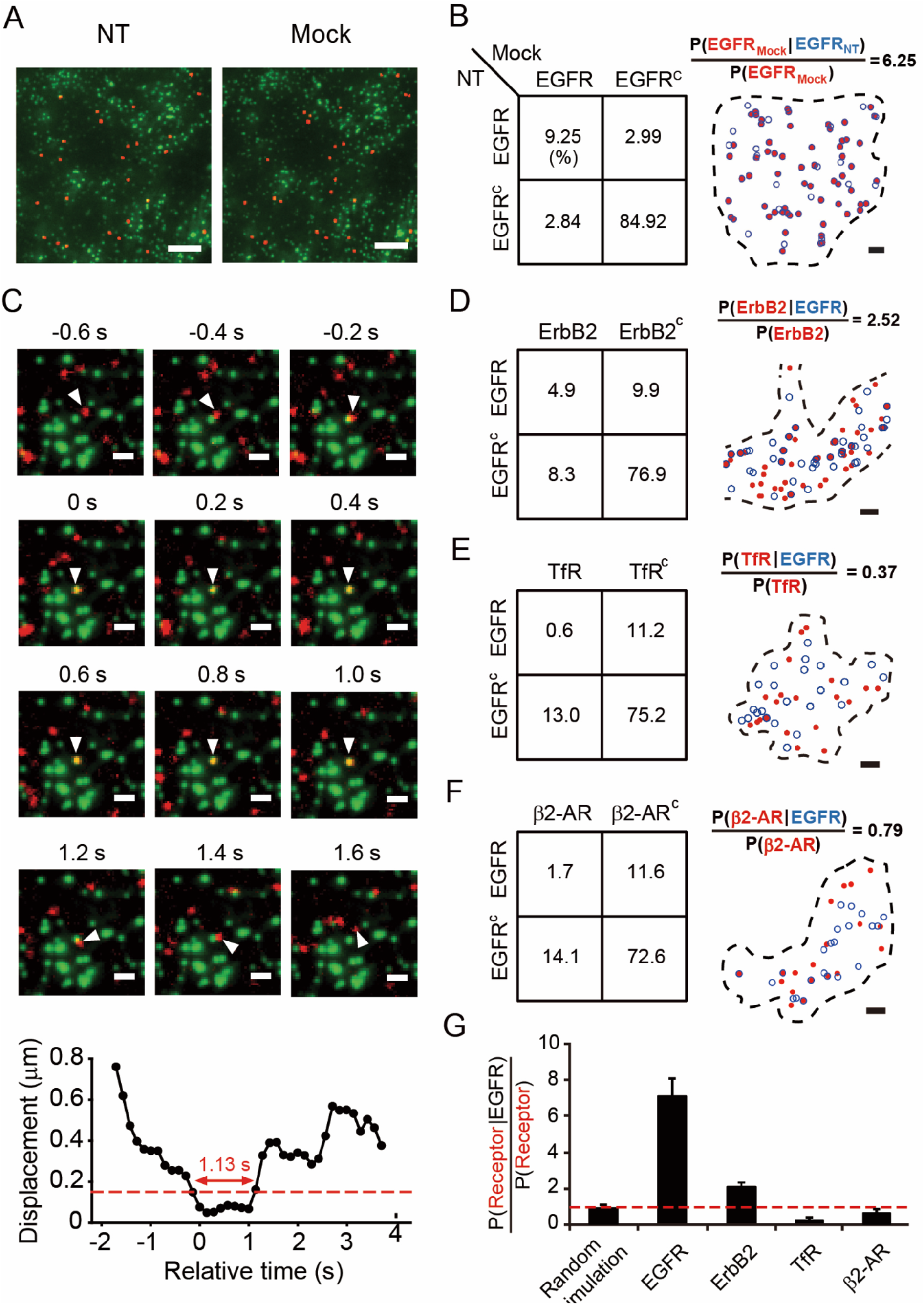
Pre-allocation property of CCPs for EGFR and the role for sorting different receptors. (A) Diffusion-based molecular colocalization analysis of CCPs with EGFR with mock treatment in a single live BSC-1 cell. (B) The quadrants of CCP ratios that contain EGFR or do not contain EGFR before and after mock treatment (*left panel*). Maps showing conditional probability of the CCP subsets containing EGFR after mock treatment, given the subset before mock treatments. Black dashed lines display cell boundaries, blue circles indicate the CCP positions before the respective treatments, and red dots indicate the CCP positions after the respective treatments (*right panel*). (C) Time-series images exhibiting the trapping process of single-molecule EGFR (red) into a single CCP (green). A white triangle in each image points to the same single-molecule EGFR undergoing the trapping process. Scale bars, 1 μm. Displacement of the single-molecule EGFR from the CCP indicated in the time-series images over time. The trapping time is determined when the displacement becomes smaller than the size of the CCP. (D, E and F) The quadrants of CCP ratios that contain or do not contain EGFR and the indicated receptors (*left panels*). The maps showing conditional probability of the CCP subset containing the indicated receptors (red dots), given the subset containing EGFR (blue circles) (*right panels*). (G) An independence test for the events between the CCP subsets containing EGFR and the indicated receptors, respectively. A red dashed line indicates the criteria of an independent event. Error bars denote s.e.m. at the single-cell level (n > 3). Scale bars, 5 μm (A, B, D, E and F).

The existence of the CCP subset pre-allocated for EGFR raises the question of whether the pre-allocation property is also applied to different receptors, which led us to additionally examine ErbB2, TfR and β2-AR using diffusion-based molecular colocalization analysis. We found the CCP subsets pre-allocated for ErbB2, TfR and β2-AR at ~12.2%, ~14.1% and ~13.8% of the total CCPs, respectively. It has been controversial whether the resistance to down-regulation of ErbB2 on the plasma membrane is due to the lack of trafficking to CCPs or rapid endosomal recycling (*24*). Our result directly shows that ErbB2 accesses CCPs without large cluster formation, accounting for why this could not be observed using conventional fluorescence microscopy. This case of ErbB2 is in contrast to the case of TfR, which constitutively traffics to CCPs by forming large clusters in the resting state.

Considering that numerous receptor types undergo CME, the ratio of the examined CCP subsets pre-allocated for each receptor was extremely high, because the sum of the subset fractions for EGFR, ErbB2, TfR and β2-AR already amounted to more than half of the total CCPs. Thus, we further explored the relationship between pre-allocated CCP subsets for different receptors. We analyzed the diffusion-based molecular colocalization of two different receptors with CCPs in a single living cell by visualizing single-molecule trajectories of the two receptors and CCPs using three-color TIRF imaging (Movie S3). Interestingly, ErbB2 tended to be loaded to the CCPs containing EGFR (Figure 3D), while TfR and β2-AR showed no such tendency (Figure 3E and 3F) in the resting state. A statistical independence test confirmed that the CCP subsets pre-allocated for EGFR and ErbB2 were partially shared, while those for EGFR and TfR, or EGFR and β2-AR were mutually exclusive (p < 0.001 and p < 0.05, respectively) (Figure 3G), suggesting the cargo-selective control of CCP pre-allocation. The previous observation that EGFR and TfR utilized different subsets after ligand stimulation might originate from their mutually exclusive pre-allocated subsets (*25*).

### PICALM as a necessary adaptor for the pre-allocated CCPs for EGFR

To better understand how a subset of CCPs is selectively pre-allocated for EGFR, we investigated the involvement of endocytic accessory proteins (EAPs) in the pre-allocation process because some EAPs were previously shown to play a key role in cargo selection and reported to be associated from the early stage of CCP formation (*14*). We screened nine different adaptors and EAPs related to EGFR (AP2μ2, FCHO, Epsin2, EPS15, EPS8, PICALM, HIP1R, GRB2 and Cbl) by quantifying the colocalization ratios between CCPs and the clusters of these endocytic adaptor proteins using two-color TIRF microscopy and particle detection analysis in the plasma membrane of living cells (Figure 4A-4J). No clustered Cbl was observed in the resting state. To verify the quantification accuracy of the colocalization, we utilized CLC-EGFP and CLC-SNAP labeled with Janelia fluorphore-549 (JF-549), a cell-permeable organic dye, which showed a colocalization ratio of ~85.4% (Figure 4A, Figure S5A, S5B and S5C). On the other hand, CLC-EGFP and Cav1-SNAP showed a colocalization ratio of ~2.9% (Figure 4B, Figure S5D, S5E and S5F) as these two components are known to be mutually exclusive (*26*).

**Figure 4.**
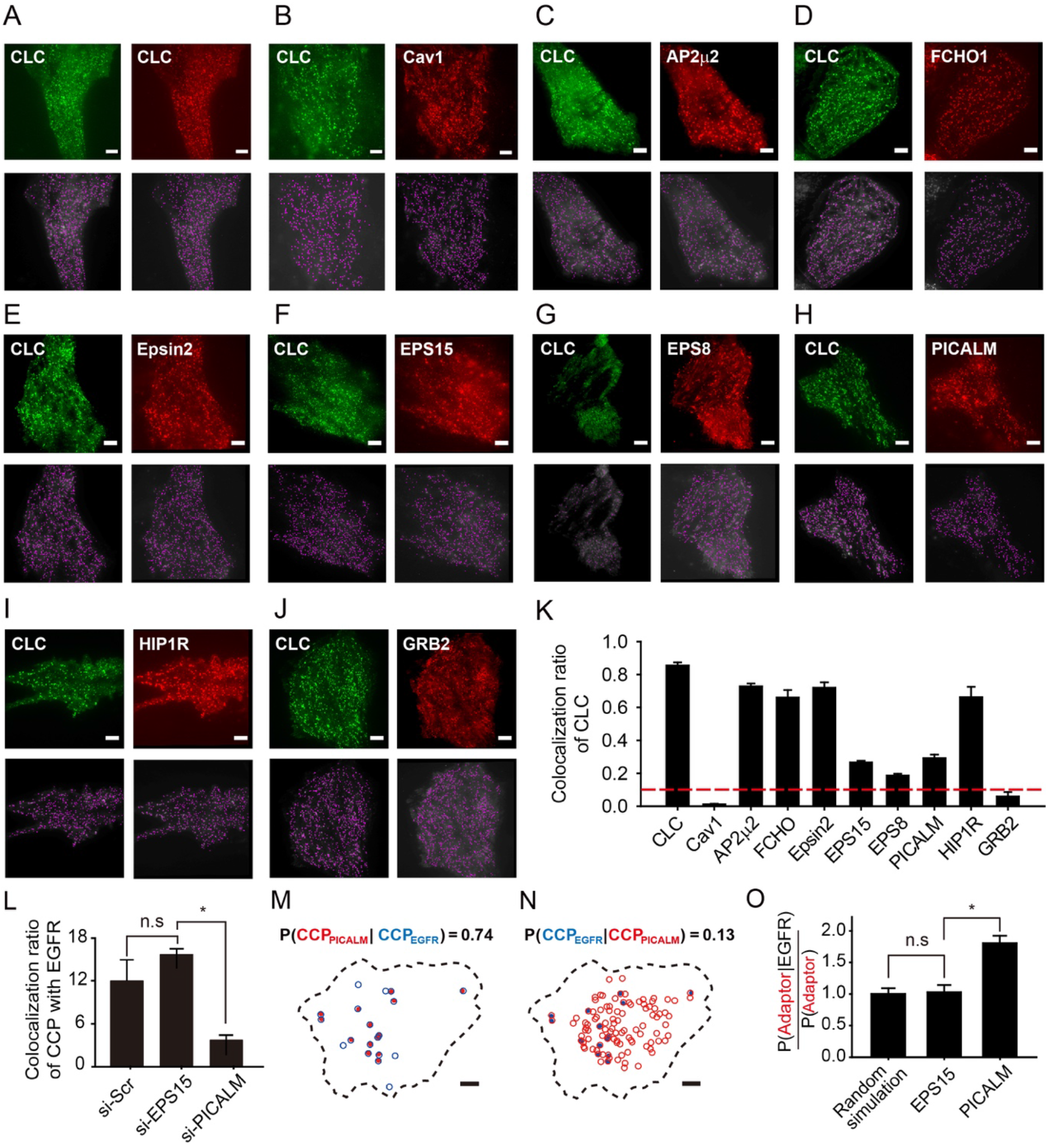
PICALM as a necessary factor for the pre-allocated subset of CCP for EGFR. (A-J) TIRFM images of EGFP-CLC and JF549-labeled SNAP-adaptors (CLC, Cav1, AP2μ2, FCHO, Epsin2, EPS15, EPS8, PICALM, HIP1R and GRB2) (*upper panels*) for the analysis of colocalization between CCPs and the clusters of each adaptor in a single living cell. The individual clusters of each adaptor objectively detected using the particle detection algorithm (magenta dots) are displayed (*lower panels*). (K) The quantitative analysis of the colocalization ratios of CCPs with the clusters of each adaptor. The colocalization ratio of CCPs with EGFRs is shown (a red dashed line). (L) A histogram for the colocalization ratio of CCPs with EGFR after the knock-down of the indicated adaptors. (M, N) Maps showing conditional probability of the CCP subsets containing PICALM (red dots), given the subset containing EGFR (blue circles) (M) or vice versa (N). (O) An independence test for the events between the CCP subsets containing EGFR and the indicated adaptors, respectively. Error bars denote s.e.m. at the single-cell level (n > 5). Scale bars, 5 μm. *p < 0.01 (Student’s *t*-test).

All adaptors and EAPs that we examined displayed a different level of colocalization with CCPs in the resting state (Figure 4K). We first searched for the adaptors or EAPs that had a sufficiently low colocalization ratio with CCPs but that was higher than the ratio of the pre-allocated CCP subset for EGFR so that they could serve an EGFR-selection role. As expected, the core components for CCP initiation, including AP2μ2 and FCHO, displayed high colocalization ratios (~72.8% and ~66.1%, respectively), which presumably did not serve an EGFR-selection role. Epsin2 and HIP1R also exhibited high colocalization ratios (~72.0% and ~66.3%, respectively), although these adaptors were plausible candidates, considering their roles in binding to the membrane, clathrin and other components. In contrast, the colocalization ratio of GRB2 (~5.9%) was lower than the ratio of the pre-allocated CCP subset for EGFR, implying that GRB2 could not be a necessary component for the pre-allocated subset. EPS15, EPS8 and PICALM were strong candidates satisfying the criteria (~26.6%, ~17.5%, and ~29.2%, respectively). However, the clusters of EPS8 completely disappeared upon Pitstop 2 treatment, implying that EPS8 was not related to the pre-allocation process, which was observed in the presence of Pitstop2.

To examine whether EPS15 and PICALM contribute to the pre-allocated CCP subset for EGFR, we knocked-down either adaptor using siRNA. EPS15 silencing exhibited no effect on the pre-allocated subset, whereas PICALM silencing significantly reduced the ratio of the subset by ~3.2-fold (Figure 4L). We directly examined the relationship between the pre-allocated CCP subset for EGFR and the CCP subset colocalized with the cluster of either adaptor by visualizing singlemolecule EGFR trajectories, CCPs, and the adaptor clusters in a single living cell using three-color TIRF imaging. Intriguingly, most of the CCPs containing EGFR were colocalized with PICALM with a conditional probability of 0.74 (Figure 4M). However, among the CCPs colocalized with PICALM, only a small portion contained EGFR with a conditional probability of 0.13 (Figure 4N). These results indicate that PICALM is a necessary but not sufficient condition for the preallocated CCP subset for EGFR. The relationship between the pre-allocated CCP subset for EGFR and the CCP subset colocalized with EPS15 was independent (Figure 4O), consistent with the siRNA result (Figure 4L).

### Facilitated EGFR Dimerization in the Pre-allocated CCPs

To investigate the role of pre-allocated CCP for EGFR, we explored the effect of EGF on the CCP subset for EGFR existing in a resting state. We quantitatively examined the changes in the molecular colocalization images between EGFRs and CCPs before and after EGF stimulus in a single living cell. Surprisingly, the fraction of the CCP subsets containing EGFR was not significantly increased by 2 nM EGF (Figure S6A), whereas the average number of EGFR molecules in a CCP was substantially elevated (~6.8-fold) (Figure S6B). We also examined the effect of EGF on the CCP subset for EGFR at the single-cell level (Figure S6C). We found that most of the CCPs in the EGFR-containing subsets with or without EGF treatment coincided, although EGF produced a small fraction of CCPs containing EGFR, which did not exist in the resting state (Figure S6D). The conditional probability that CCPs contained EGFR before EGF treatment, given the CCP subset containing EGFR after EGF treatment, was 0.71 (Figure S6E). This observation showing that the subset of CCP pre-allocated for EGFR was primarily utilized after EGF stimulation implied that the role of the pre-allocated CCPs might be related to the receptor activation. We tested this hypothesis by examining the dimerization of EGFR inside the pre-allocated CCPs using bimolecular fluorescence complementation (BiFC) of mEos3.2 (27). We observed ~7.1% of the pre-allocated CCPs containing EGFR dimers in a resting state. Interestingly, this portion was significantly decreased by Pitstop2 treatment, whereas the portion is increased by Dynasore treatment (Figure 5A). Considering the role of these inhibitors on the CCP development, these results suggest that the pre-allocated CCPs for EGFR at the open-neck stage presort EGFR on the plasma membrane, which is subsequently dimerized as the neck of the CCPs is sealed by dynamin (Figure 5B).

**Figure 5.**
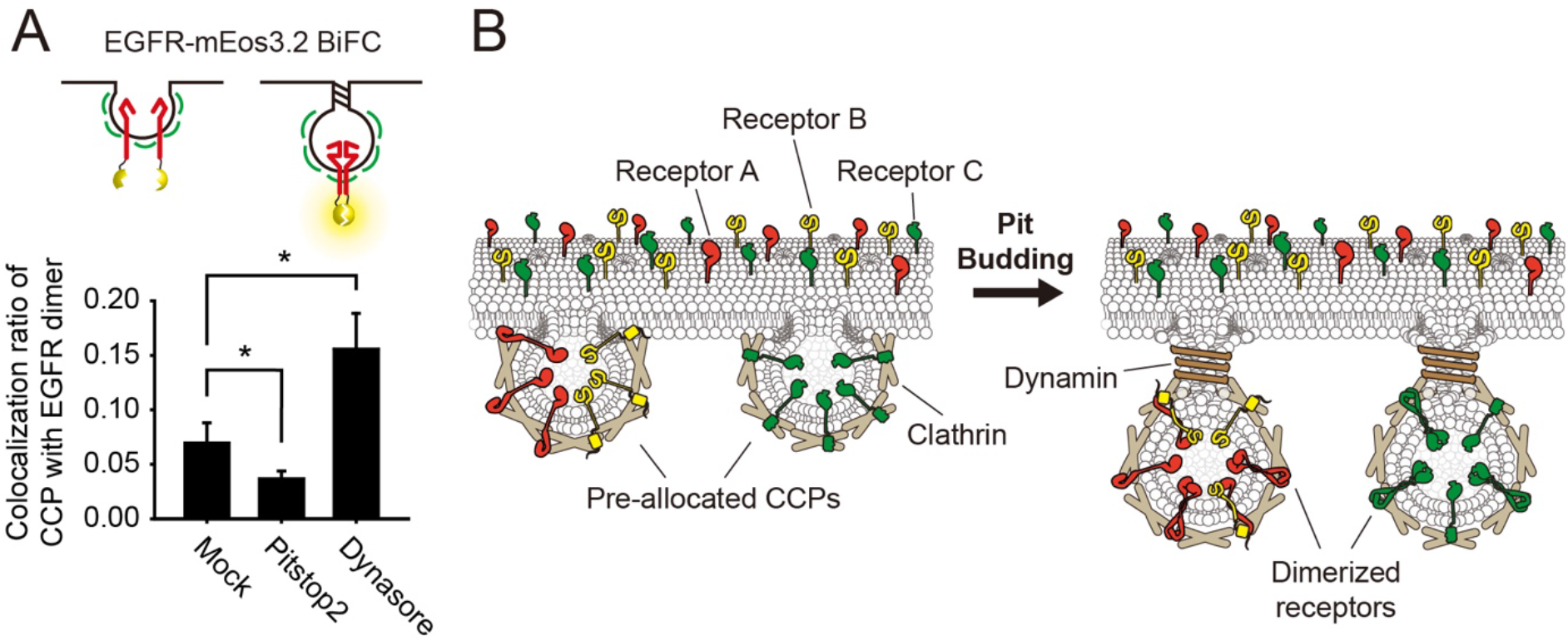
The role of CCP pre-allocation. (A) A histogram for the colocalization ratio of CCP with dimerized EGFR visualized utilizing the bifunctional fluorescence complementation (BiFC) of EGFR-mEos3.2 after Pitstop 2 or Dynasore treatment. Error bars denote s.e.m. at the single-cell level (n > 3). *p < 0.01 (Student’s *t*-test) (B) The illustration of the pre-allocation property of CCPs that different types of receptors are sorted in the shared or exclusive subsets of CCPs in the resting state and dimerized upon the recruitment of dynamin.

## Discussion

We observed the pre-allocation process of CCPs for receptors in the resting state by utilizing the diffusion-based molecular colocalization analysis. The involvement of EGFR internalization pathways is considered to be initiated upon EGFR activation (*17*). However, the existence of a CCP subset pre-allocated for EGFR showed that CME is involved in EGFR signaling prior to EGFR activation. This early involvement of the internalization pathway in EGFR signaling was limited to CME as the pre-allocation event was not observed for caveolae (Figure S7), which serve the clathrin-independent endocytosis (CIE) pathway.

The role of CCP pre-allocation for EGFR seemed to coordinate EGFR signaling prior to its activation by ligands. The CCP subset pre-allocated for EGFR was primarily utilized after the EGF stimulus, and the subset was not affected by the adaptors involved after the EGF stimulus, including Epsin2, EPS8, EPS 15, Grb2, and Cbl (*28–32*). Furthermore, EGFR dimerization was markedly elevated inside the pre-allocated CCP subset after dynamin recruitment in the resting state. EGFR dimerization at the CCPs might be derived from an increase in the local concentration generated as the neck of CCPs is closed upon dynamin recruitment, which substantially reduces the effective volume of CCPs by isolating their connectivity with the plasma membrane (33–35) (Figure 5B). This model may account for a key feature typifying CCPs, a small-sized uniform structure. The recent finding that EGFR signaling is not affected by the disassembly of clathrin plaques that form a large flat region further supports this model (*36*). Although it is not clear whether causality exists between EGFR dimerization at CCPs and phosphoinositide conversion at CCPs, it is remarkably coincident that both changes inside CCPs appeared after dynamin recruitment (*37*). Considering that the phosphoinositide compositions in CCPs are converted to transport cargo proteins from the plasma membrane to an endosome, the function of EGFR dimerization in CCPs might be closely related to the endosomal signaling of EGFR.

It remains a puzzle how pre-allocated CCPs are generated. The fates of individual CCPs to load different cargos are spatially distinct after the maturation stage (Figure 3), yet the essential factors involved at the initiation stage are common: AP2, PI(4,5)P2, clathrin, FCHo1/2 (*14*). Cargos might play a role to differentiate each CCP in addition to control the maturation time of the CCP (*15, 38*) by interacting with AP2 after the initiation of the pit because AP2 is capable to interact with the cargos and the cargos themselves are the distinct factors from the common EAPs. This hypothesis is consistent with the data that the fraction of the pre-allocated subset for EGFRΔAP2 was significantly reduced (Figure 2D). Although EGFR requires PICALM to be pre-allocated in CCPs, PICALM alone was not sufficient to hold EGFR in CCPs (Figure 4). Together with PICALM, other multiple co-adaptors might be additionally required to completely generate the selectivity of CCP pre-allocation for EGFR (*12*).

Our observations suggest that CCPs act as shredded organelles that sort receptors on a crowded plasma membrane prior to ligand-induced receptor activation (Figure 5B). The pre-allocation process of CCPs provides crucial information on the mechanism of how CCPs control not only the competition among receptors by intrinsically limiting their capacity for one type of a receptor but also the cooperation among receptors by gathering functionally correlated receptors prior to receptor activation.

## Materials and Methods

### Reagents

Mouse monoclonal antibody against EGFR (#MS-396) was purchased from Thermo Fisher Scientific (Barrington, IL). Alexa 647-conjugated secondary antibodies for immunofluorescence were purchased from Thermo Fisher Scientific (Barrington, IL). EGF (#E9644) was purchased from Sigma Aldrich (St. Louis, MO). The cell-permeable clathrin-mediated endocytosis inhibitor Pitstop2 (#ab120687) was purchased from Abcam (Cambridge, MA). 1-BtOH (#B0713) and t-BtOH (#36,053-8) were purchased from Samcheon (Seoul, Korea) and Aldrich Chemical Company (St. Louis, MO), respectively. Cy3b NHS ester was reacted with BG-NH2 (S9148S, New England Biolabs) in DMF with shaking at 30°C overnight according to the manufacturer’s instructions. The solvent was evaporated under vacuum, and the product was dissolved in distilled water after purification by HPLC. Cell-permeable Halo ligand-labeled dye, Halo JF549, was a gift from Dr. Luke D. Lavis.

### Plasmid DNA

The EGFP-conjugated CLC plasmid was constructed by subcloning EGFP into the pcDNA3.1 vector to generate pcDNA3.1/EGFP-His from the pEGFP-N1 vector (#6085-1, Clonetech, Mountain View, CA). Then, CLC was subcloned into pcDNA3.1/EGFP at the N-terminus of each tag with primers as follows: 5’-GCTCTAGAAT GGT GAGCAAGGGCGAGGAGCTGTT and 5’-GATCCACCGGTCTTGTACAGCTCGTCCATGCCGAGAGTG. To construct mEos3.2-tagged protein at the N-terminus of EGFR, we subcloned human EGFR into the pcDNA3.1 vector (V800-20, Invitrogen) with the following primers: 5’-CGCAAATGGGCGGTAGGCGTG and 5’-CCGCGGTTGGCGCGCCAGCCCGACTCGCCGGGCAGAG, and 5’-GGCGCGCCAACCGCGGCTGGAGGAAAAGAAAGTTTGC and 5’-AGCTTTGTTTAAACTTATGCTCCAATAAATTCACTGCT. Then, mEos3.2 excised from pEGFP-N1/mEos3.2, a kind gift from Dr. Tao Xu (Chinese Academy of Science), was inserted into the site between the signal and mature peptides of EGFR with the following primers: 5’-GGCGCGCCACATCATCACCATCACCATATGAGTGCGATTAAGCCAGA C and 5’-TCCCCGCGGCCCTCCACTCCCACTTCGTCTGGCATTGTCAGGCAA. To construct mEOS3.2-tagged EGFR ΔAP2, we generated Y974A, L1010A and A1011A mutant from mEOS3.2-EGFR using overlap PCR with the following primers: 5’-ATCGATGTCTACATGATCATGGTCAAG, 5’-CATCAGGGCACGGCGGAAGTTGGAGTC and 5’-GACTCCAACTTCGCCCGTGCCCTGATG and 5’-GAGCTCAGGAGGGGAGTCCGTGACGTG. To construct SNAP-tagged EGFR, the SNAP-tag gene from the pSNAPf vector (N9183S, New England Biolabs) was subcloned into the pcDNA3.1/mEos3.2-EGFR gene with the following primers: 5’-GGCGCGCCACAT CAT CACCAT CACCAT AT GGACAAAGACT GCGAAAT G and 5’-TCCCCGCGGCCCTCCACTCCCACTACCCAGCCCAGGCTTGCCCAG. To construct SNAP-tagged ErbB2, we subcloned ErbB2 into pcDNA3.1/SNAP-EGFR using overlap PCR with the following primers: 5’-CGCAAATGGGCGGTAGGCGTG, 5’-CAGCTCCATCCCGCGGCCCTCCACT and 5’-AGT GGAGGGCCGCGGAT GGAGCT G, 5’-AGCTTTGTTTAAACTCACACTGGCAGTCCAGACCCAG. To construct SNAP-tagged TfR, we subcloned TfR into pcDNA3.1/SNAP-EGFR with the following primers: 5’-TCCCCGCGGATGGATCAAGCTAGATCA and 5’-AGCTTTGTTTAAACTAAAAACTCATTGTCAATGTC. The corresponding templates were obtained from Taylor MJ (Addgene plasmid #27680 for CLC), Matthew Meyerson (Addgene plasmid #11011 for EGFR), Alexander Sorkin for EGFR ΔAP2, and David Perrais (Addgene plasmid #61547 for TfR).

The Halo-conjugated GRB2 and PICALM were constructed by subcloning (1) Halo into the pcDNA3.1 vector (#V800-20, Invitrogen, Carlsbad, CA) to construct pcDNA3.1/Halo-His from pENTR4-Halo obtained from Eric Campeau (Addgene plasmid #29644) with the primers 5’-GCTCT AGAGGAGGGAT GGCAGAAAT CGGT ACT GGC and 5’-GATCCACCGGTGCCGGAAATCTCGAGCGTCGA and (2) GRB2 obtained from Dominic Esposito (Addgene plasmid #70383) and PICALM, a kind gift from Jin-moo Lee, were subcloned into the C-terminal of the pcDNA3.1/Halo-His vector with the following primers: 5’-GGAATTCATGGAAGCCATCGCCAAATATGACTTCAAA and 5’-GCTCTAGAGACGTTCCGGTTCACGGGGGT for GRB2 and 5’-CGGGATCCATGTCTGGCCAGAGCCTGACG and 5’-ATAAGAATGCGGCCGCTTCATAAACTGTATCTGTGCTCC for PICALM. Eps15-mCherry (Addgene plasmid #27696), AP2μ2-mCherry (Addgene plasmid #27672), FCHo1-mCherry (Addgene plasmid #27690), Epsin2-mCherry (Addgene plasmid #27673), Eps8-mCherry (Addgene plasmid # 29779), and Hip1R-tDimerRFP (Addgene plasmid #27700) were a gift from Christien Merrifield.

### Cell handling

BSC1 cells were grown at 37°C and 5% CO2 in EMEM (#M5650, Sigma Aldrich, St. Louis, MO) supplemented with 10% fetal bovine serum, 100 U/ml penicillin, 100 μg/ml streptomycin and 0.25 μg/ml amphotericin B. BSC1 cells (1 × 10^5^) were placed on collagen-coated coverslips and imaged 12 to 18 h after plating. In the experiments designed to evaluate endogenous EGFR labeled by anti-EGFR Fab-Alexa 647, CLC conjugated with an EGFP stable cell line was placed on culture well-attached coverslips. For the experiment using BSC1 cells transiently transfected with SNAP-tagged EGFR, TfR and ErbB2, transfected BSC1 cells were labeled with BG-Cy3b, prior to placing on the coverslips. For the experiment using BSC1 cells transiently transfected with Halo-tagged PICALM and GRB2, transfected BSC1 cells were labeled with Halo-JF549, prior to placing on the coverslips.

### Stable cell line generation

BSC1 cells were transfected to express EGFP-conjugated CLC by Lipofectamine LTX (#15338100, Thermo Fisher Scientific, Barrington, IL) according to the manufacturer’s instructions. For selection of pcDNA3.1/EGFP-CLC-expressing BSC1 cells, 1 mg/ml geneticin antibiotic (#10131035, Thermo Fisher Scientific, Barrington, IL) was added to the growth media 2 days after transfection. Then, pcDNA3.1/EGFP-CLC low-expression BSC1 cells were sorted by the FACS machine (MoFlo™, Beckman Coulter, Brea, CA) after a 3-day incubation in the presence of antibiotic. Homogenous pcDNA3.1/EGFP-CLC stably expressing BSC1 cells were obtained after the sorting process was repeated three times.

### Plasmid DNA transfection

Transient co-expression of SNAP-EGFR, EGFR ΔAP2, ErbB2 and TfR with EGFP-CLC was achieved by transfection with Lipofectamine LTX according to the manufacturer’s procedure. For 12-well plates, 1.8 μg of plasmid DNA encoding SNAP-EGFR, EGFR ΔAP2, ErbB2, or TfR and 0.2 μg of plasmid DNA encoding EGFP-CLC were diluted in 50 μl of Opti-MEM (#31985-070, Thermo Fisher Scientific, Barrington, IL) containing 2 μl of plus reagent (#11514-015) and combined for 5 min at room temperature with a solution of 5 μl of Lipofectamine LTX diluted in 50 μl of Opti-MEM. Cells were imaged 48 h after transfection.

For transient co-expression of EGFP-CLC with adaptor plasmids, the following ratios of EGFP-CLC to adaptor plasmid DNA were used for 12-well plates to achieve similar expression levels: Halo-GRB2 (2:1), PICALM (1:2), Eps15-mCherry (4:1), AP2μ2-mCherry (1:4), FCHo1-mCherry (4:1), Epsin2-mCherry (4:1), Eps8-mCherry (1:1), and Hip1R-tDimerRFP (4:1). In total, 1 μg of DNA was transfected with Lipofectamine LTX by the same procedure described above.

### Fluorescently labeled Fab fragment of antibody preparation

Neutral mouse monoclonal antibody against EGFR (#MS-396, Thermo Fisher Scientific, Barrington, IL) was cleaved into Fab and Fc fragments and purified using a Pierce™ Fab preparation kit (#44985, Thermo Fisher Scientific, Barrington, IL) according to the manufacturer’s protocol. After purification, the Fab fraction was concentrated for a better labeling yield using 10K centrifugal filter units (#UFC501024, Millipore, Billerica, MA). Then, Fab was labeled with Alexa 647 using an Alexa^®^ Fluor 647 antibody labeling kit (#A-20186, Thermo Fisher Scientific, Barrington, IL), as suggested in the manufacturer’s protocol. After labeling, anti-EGFR Fab-Alexa647 was concentrated to establish an experimental working concentration (200 nM).

### Glass coverslip preparation

Glass coverslips (#0111580, Marienfeld Laboratory Glassware, Lauda-Konigshofen, Germany) were cleaned using a water bath sonicator at 42°C for 30 min, first in acetone and then for 30 min in 1% hydrofluoric acid. Then, coverslips were washed with Milli-Q^®^ (Millipore, Billerica, MA)-filtered water. Next, the coverslips were sterilized by soaking in 100% ethanol under UV light for more than 30 min and washed three times with PBS. The coverslips were coated with collagen (20 μg/ml) dissolved in Milli-Q-filtered water containing 0.5 M acetic acid for 1 h. Prior to seeding the cells, coverslips were washed 3 times with PBS. One silicon well of 3-to 10-μl-sized Grace Bio-Labs CultureWell chambered cover glass (#GBL103350, Sigma Aldrich, St. Louis, MO) was attached on the center of the washed coverslip before collagen coating.

### Live cell imaging

Cells prepared for each experiment were placed on coverslips, as described in the cell handling methods. Then, the coverslips were placed in the live cell chamber after washing three times with PBS to minimize auto-fluorescence from phenol red media and imaged in phenol red-free media containing 2.5 mM protocatechuic acid (PCA, #03930590, Sigma Aldrich, St. Louis, MO), 0.2 U/ml protocatechuate 3,4-dioxygenase (PCD, #P8279, Sigma Aldrich, St. Louis, MO) and 1 mM β-mercaptoethylamine (MEA, #M9768, Sigma Aldrich, St. Louis, MO) imaging media.

For anti-EGFR Fab-Alexa 647 labeling, cells were pre-incubated with phenol red-free media containing 200 nM anti-EGFR Fab-Alexa 647 at 37°C for 15 min. For Pitstop 2 treatment experiments, cells were serum-starved for 4 h and preincubated for 15 min with 30 μM Pitstop 2. For three-color imaging of CLC, adaptor, and EGFR, we first identified an appropriate cell expressing CLC and adaptor and then labeled it with anti-EGFR Fab-Alexa 647. Then, two-color imaging of EGFP-CLC and EGFR in the 488-647 nm channel in Pitstop 2-containing imaging media was performed as mentioned above, followed by imaging of EGFP-CLC and an adaptor in the 488-561 nm channel. For sequential imaging of SNAP-EGFR/TfR/ErbB2 and anti-EGFR Fab-Alexa 647 with EGFP-CLC, we first imaged EGFP-CLC and EGFR labeled with anti-EGFR Fab-Alexa 647 in PCA, PCD, and MEA-containing imaging media with Pitstop 2 treatment in the 488-647 nm channel and then imaged EGFP-CLC and SNAP-EGFR/TfR/ErbB2 in the 488-561 nm channel as mentioned above.

### Acquisition and analysis of image data

Imaging was performed using an objective-type total internal reflection fluorescence microscope (TIRFM). During the entire course of imaging, samples were maintained in a live-cell imaging chamber (Chamlide TC-A, Live Cell Instrument). An EGFP protein was excited using a wavelength-tunable ion laser (543-BS-A03, Melles Griot) with an intensity of 20 to 100 mW/cm^2^. An mEos3.2 protein was excited using a 561-nm laser (YLK 6150T, Lasos) with an intensity of 20 to 30 W/cm^2^ and photoactivated using a 405-nm laser (DL-450-120, Crystal Laser) for 100 to 1,000 ms, depending on the expression level with an intensity of 0. 2 to 0.5 W/cm^2^. An Alexa Fluor 647 dye was excited using a 642-nm laser (2RU-VFL-P-1000-642, MPB Communications) with an intensity of 20 to 100 W/cm^2^. All instrument operations and data acquisition were controlled using MetaMorph (Molecular Devices) and custom plug-ins written in MATLAB (The MathWorks). Multiple particle tracking was performed based on U-track (*39*) as previously described (*40*). The diffusion coefficient from a single trajectory was calculated from the MSD with the following two-dimensional free diffusion model equation:

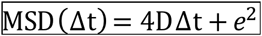

where 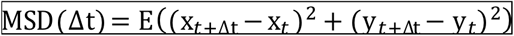 and 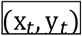 are the Cartesian coordinates of a particle at the t^th^ point of its trajectory and *e* is a localization error. Trajectories with a duration longer than ten frames were used to calculate the diffusion coefficient using the four time lags of MSD.

For detecting CCP locations, 40 frames of CLC-EGFP images were averaged to match the diffusivity of CCP and imaging acquisition speed, which allows to use a low power of the excitation laser to minimize photobleaching and significantly enhances a signal-to-noise ratio. The CCP and EGFR positions were colocalized after drift correction and dual-channel alignment of the EGFR trajectories. Immobilized EGFR within the size of CCPs (typically, 50 to 120 nm) from the CCP location in the same time-frame was considered trapped, and the number of trapped EGFRs in each CCP was calculated for the analysis.

Translational drift during cellular imaging was corrected by calculating an affine transformation matrix between every consecutive images. The locations of CLC clusters or 100-nm diameter gold particles were used for the correction marker. When using CLC clusters, the matrix was obtained using the eight-point algorithm with the random sample consensus (RANSAC) algorithm fitted to the marker data. When using gold particles, the particles were added before imaging, and cells near the particles bound to glass were used. For the registration of images from two EM-CCDs, microspheres (Crimson) were imaged with a dual-color channel after every imaging experiments. The transformation matrix was obtained as mentioned in the drift correction method using bead images.

### Simulation generation

Simulated data of EGFR and CCP were generated by making random diffusive and immobilized particles. One hundred thousand EGFR particles diffused freely with 0.2 μm^2^/s in a 512×512 pixel (1 pixel = 100 nm) domain with 1,000 CCPs of a 100-nm radius distributed at random locations. As the EGFR particle and CCP met, EGFR had a chance, *p_trap,* to be trapped in the CCP, which could trap a maximum of 20 EGFR in each cargo, and the probability, *p_off*, determined how fast EGFR escaped the CCP. For the random trapping model, every CCP had same *p_trap* and *p_off*, while for the predetermined CCP model, *p_trap* was greater for the predetermined ones. In every frame, only a few hundred EGFR particles were designated to be shown, with an average of 25 frames, and each location showing EGFR and CCP in every frame was drawn with an appropriate point spread function in the image for comparison with the real data.

## Supporting information

## Author contributions

Conceptualization, D.K. and Y.K.; Methodology, D.K. and Y.K; Software, D.K. and H.A.; Investigation, Y.K., D.K., H.A., K.Z., M.J., S.P., and Y.C.; Writing – Original Draft, D.K. and Y.K.; Writing – Review & Editing, D.K., Y.K., and S.R.; Funding Acquisition, S.R. and D.K.; Resources, N.L.; Supervision, S.R.

## Data and materials availability

The authors declare that the data supporting the findings of this study are available within the paper and its supplementary information files.

## Supplementary Materials

Figure S1. Related to Figure 1. Diffusion-based molecular colocalization between EGFR and CCP

Figure S2. Related to Figure 1. Diffusion-based molecular colocalization criteria determination

Figure S3. Related to Figure 2. Validation of specificity of the diffusion-based molecular colocalization of EGFR with CCP

Figure S4. Related to Figure 3. Pre-allocation property of CCPs for EGFR

Figure S5. Related to Figure 4. PICALM as a necessary factor for the preallocated subset of CCP for EGFR

Figure S6. Related to Discussion. Cav1 colocalization with EGFR in live mEGFP-Cav1 expressing BSC1 cell line

Movie S1. Related to Figure 1. Diffusion-based molecular colocalization between CCSs and single-molecule EGFRs

Movie S2. Related to Figure 3. Transient trapping of single-molecule EGFR into a CCP in a resting state

Movie S3. Related to Figure 5. Three-color molecular colocalization among CCPs, single-molecule TfR, and single-molecule EGFR

## Supplementary Materials

**Figure S1.**
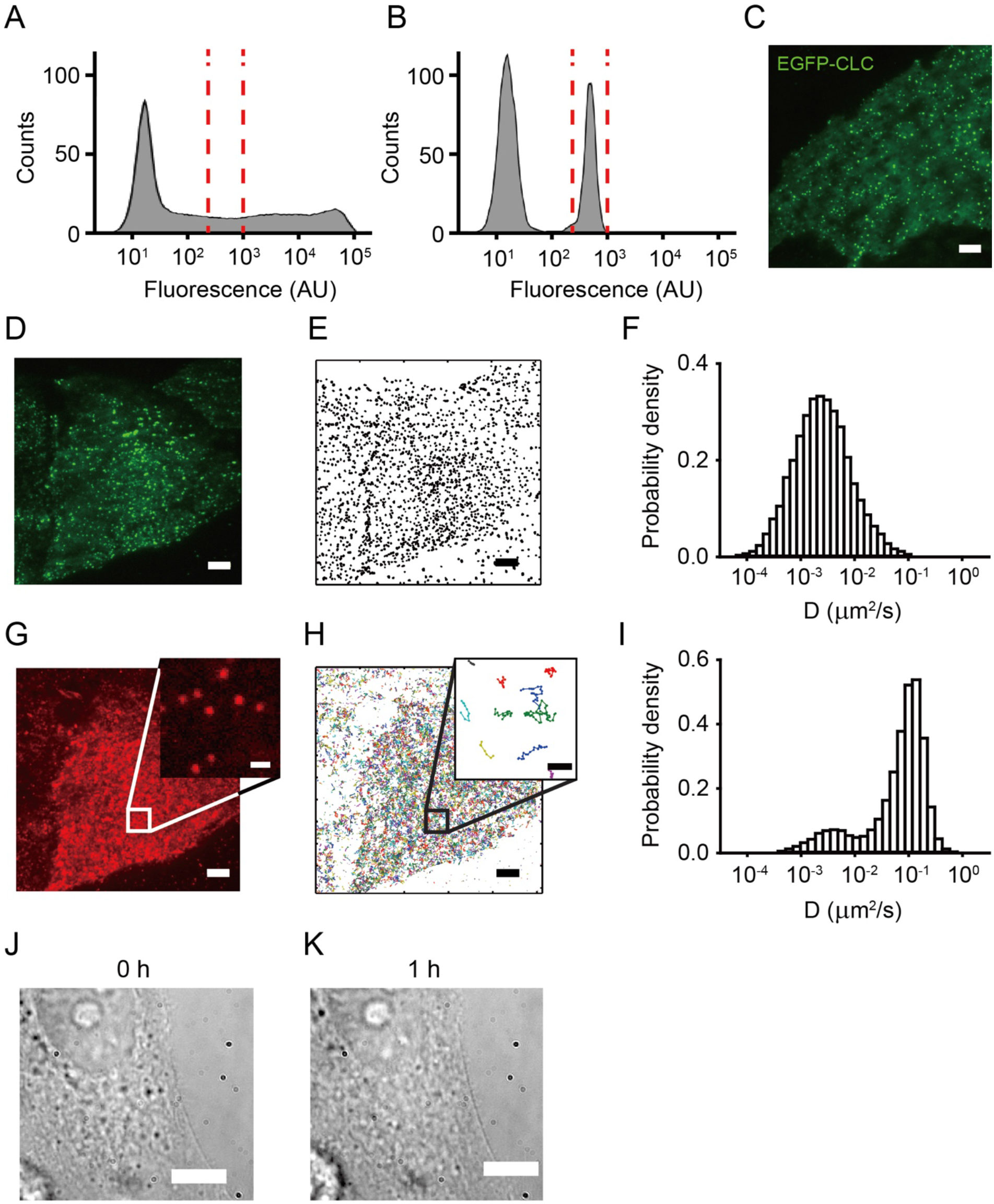
Related to Figure 1. Diffusion-based molecular colocalization between EGFR and CCP. (A-C) EGFP-CLC stable cell line generation. Flow cytometry histogram shows EGFP-CLC transiently transfected in the BSC1 cell line (A), the region delimited by vertical lines indicates the sorted population and EGFP-CLC stably expressed in the BSC1 cell line after antibiotic-based selection and low-expression population sorting from EGFP-CLC transiently transfected in the BSC1 cell line (B). (C) TIRFM image shows normal formation and distribution of CCPs on the plasma membrane of the EGFP-CLC stably expressed in BSC1 cell line. (D-I) Simultaneous two-color single-particle tracking analysis for CCP and EGFR. TIRFM images show distributions of CCP (D) and EGFR (G) in Alexa 647-conjugated Fab fragment of anti-EGFR antibody labeled EGFP-CLC stably expressed in BSC1 cell line; inset image shows a single frame of single-molecule level Alexa 647-conjugated Fab fragment-labeled EGFR from the boxed region of (G). Map of accumulated CCP (E) and EGFR (H) trajectories acquired for approximately 5 min in a single BSC1 cell; 5,000 trajectories are shown, and only 10 randomly chosen trajectories are shown in inset from (H). Histograms show that diffusion coefficient distributions of CCP (F) and EGFR (I). (J, K) Cell viability after imaging. DIC images were taken before (J) and after (K) diffusion-based single-molecule colocalization imaging, with 15-min 405-, 488- and 647-nm illuminations in a single EGFP-CLC stably expressed in a BSC1 cell. Scale bars, 5 μm (C, D, E, G, H, J and K) and 1 μm (the inset of G and H).

**Figure S2.**
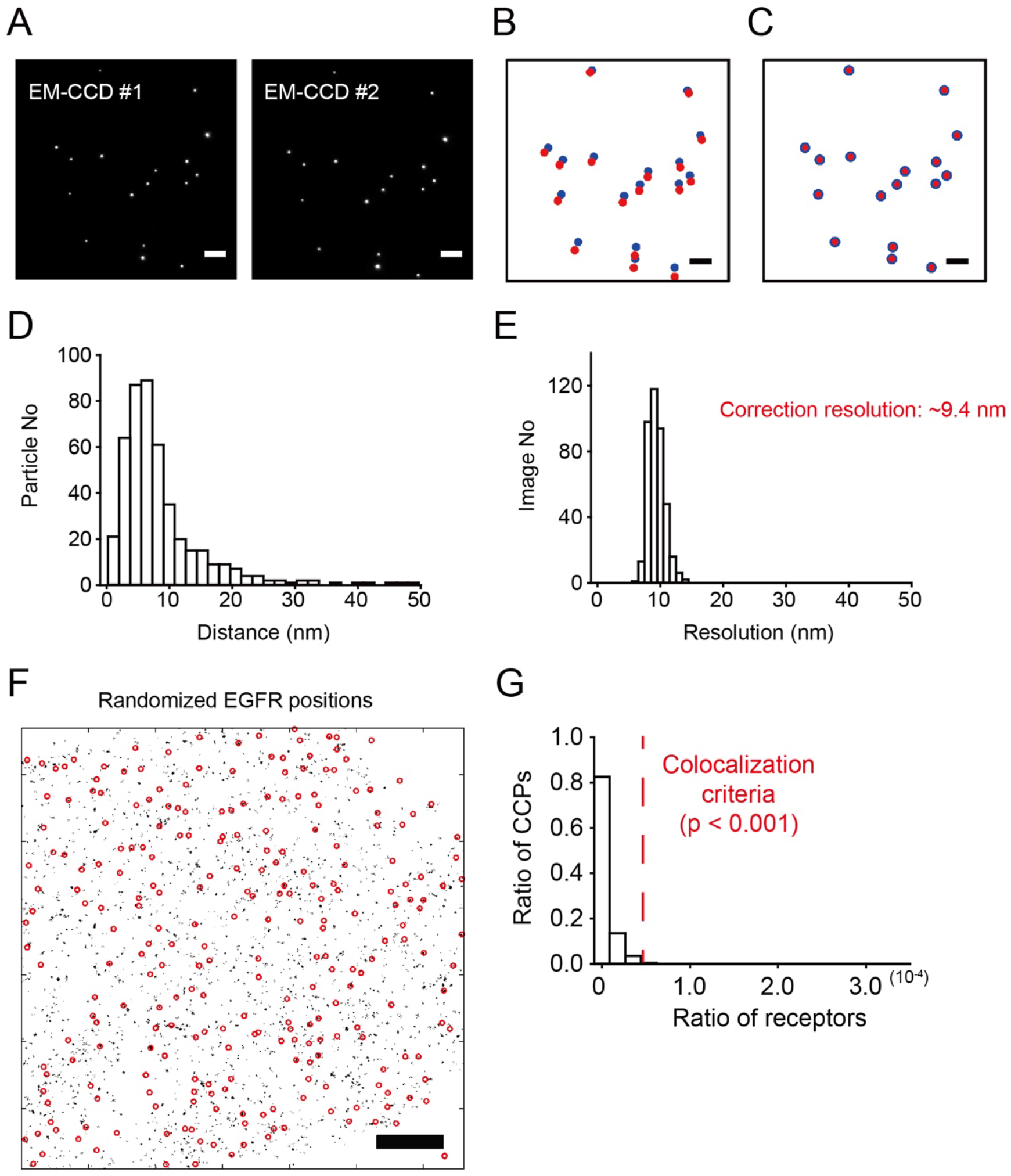
Related to Figure 1. Diffusion-based molecular colocalization criteria determination. (A-E) Two-color image registration by bead images from two different EM-CCDs. (A) TIRFM images show simultaneous snapshots of the beads from two different EM-CCDs. Plot shows a merge of coordination of beads before (B) and after (C) affine transformation from two different EM-CCDs, blue and red dots indicate coordination of beads from EM-CCD#1 and EM-CCD#2, respectively. Histograms show the distributions of distances between coordination of beads from two different EM-CCDs after affine transformation from single (D) and multiple (E) bead images. (F, G) Molecular colocalization between CCP and EGFR by diffusion-based single-molecule colocalization. (F) Overlaid map of accumulated CCPs (red circle) and randomized EGFRs (black dot), which were randomly redistributed from the cell in Figure 1, (G) A histogram represents the distribution of the ratio of CCP with respect to the ratio of randomized EGFR. Scale bars, 5 μm.

**Figure S3.**
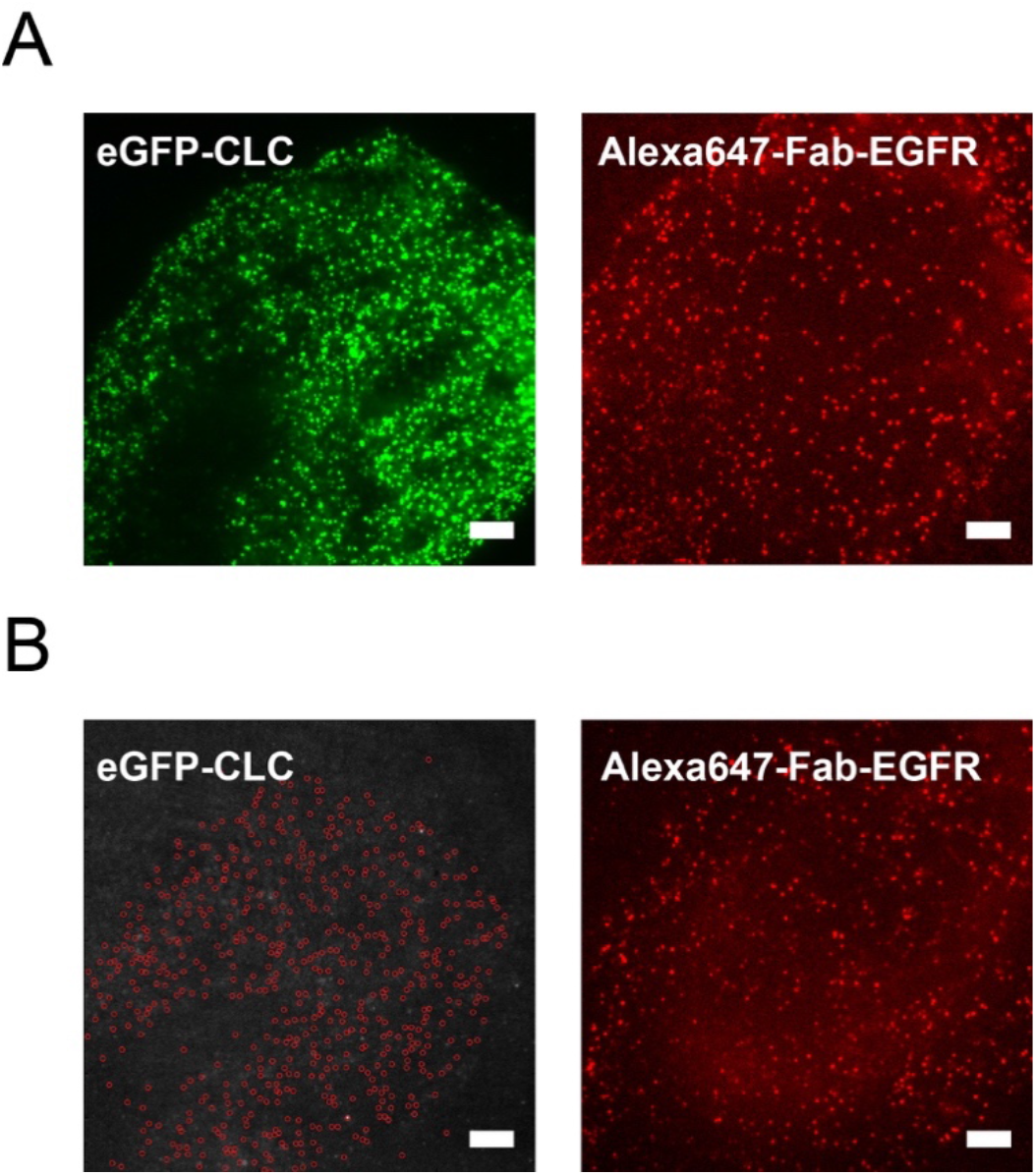
Related to Figure 2. Validation of specificity of the diffusion-based molecular colocalization of EGFR with CCP. TIRFM images show the EGFP-CLC (*left panels*) and Alexa 647-conjugated anti-EGFR antibody Fab fragment-labeled EGFR (*right panels*) before (A) and after (B) acute treatment with 2% 1-Butanol. Red circles indicate the coordination of CCPs, which were located before 1-BtOH treatment. Scale bars, 5 μm.

**Figure S4.**
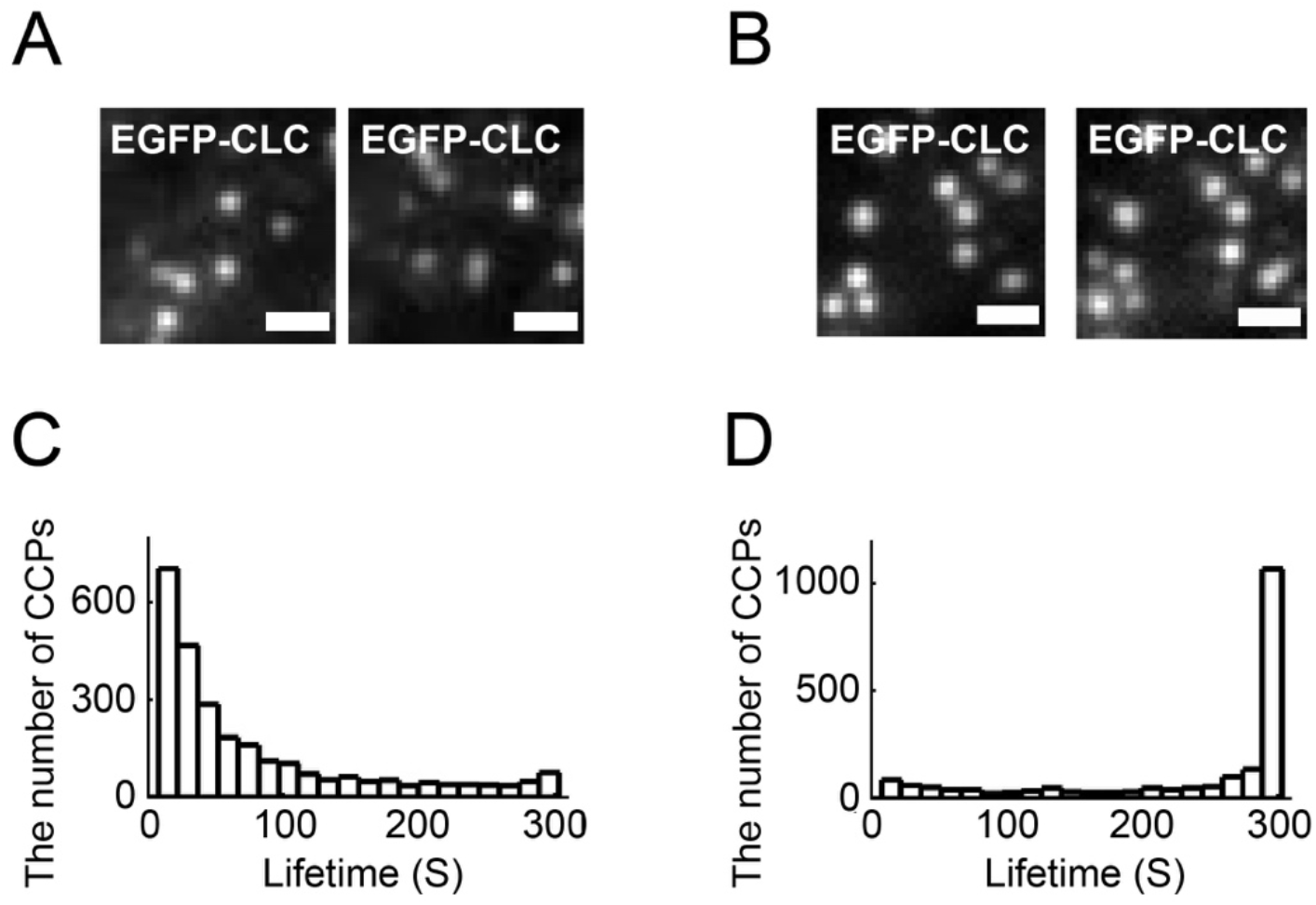
Related to Figure 3. Pre-allocation property of CCPs for EGFR. TIRFM images show that distribution of CCPs at the initial 0 sec (*left panels*) and the 300 sec (*right panels*) time-points in EGFP-CLC stably expressed in BSC1 cell line imaging without (A) and with (B) Pitstop 2 pre-incubation. Scale bar, 1 μm. Histograms show the distributions of lifetime of CCPs without (C) and with (D) pre-incubation of Pitstop 2.

**Figure S5.**
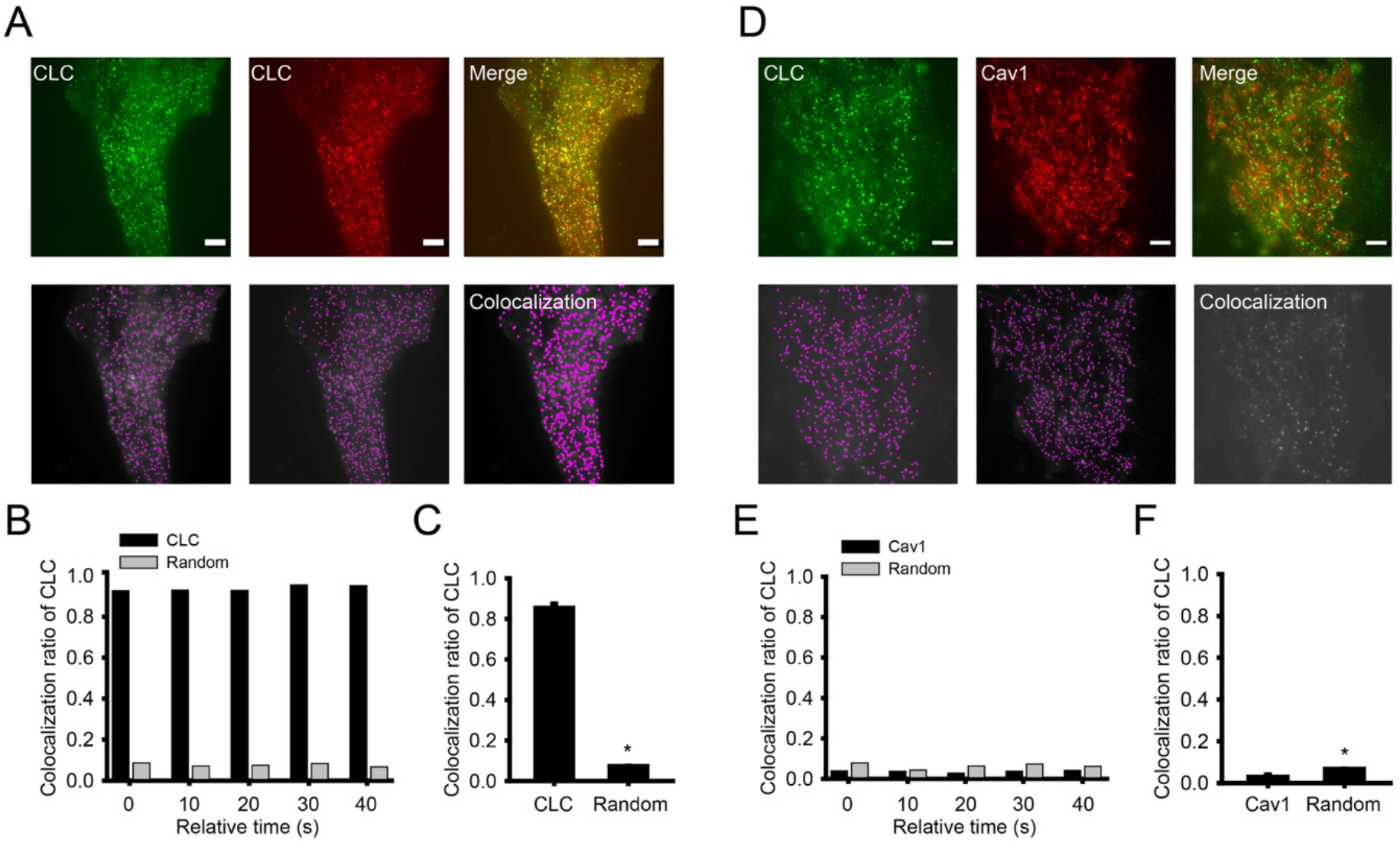
Related to Figure 4. PICALM as a necessary factor for the preallocated subset of CCP for EGFR. TIRFM images (*upper panels*) show the distributions of CCPs (*left*), indicated EAPs (*middle*) and merge (*right*) and dot detection analyzed images (*lower panels*) of CCPs (*left),* indicated EAPs (*middle*) and colocalization (*right*) in EGFP-CLC and indicated EAPs transiently co-expressed in BSC1 cell line. (A) CLC, (D) Cav1. The magenta dot indicates the CCPs or EAPs, detected by dot detection analysis as a single dot from TIRFM image, Scale bars, 5 μm. Histograms represent co-localization ratio of JF549-SNAP-CLC with EGFP-CLC (B, black bar) or mEGFP-Cav1 (E, black bar) and colocalization of randomized JF549-SNAP-CLC with EGFP-CLC (B, grey bar) or with mEGFP-Cav1 (E, grey bar) at different time points in a single living cell. Histograms show multiple cell colocalization ratio of JF549-SNAP-CLC and randomized JF549-SNAP-CLC with EGFP-CLC (C) and mEGFP-Cav1 (F). Error bars for denote s.e.m. (n > 8). Scale bars, 5μm. *p < 0.01 (Student’s *t*-test).

**Figure S6.**
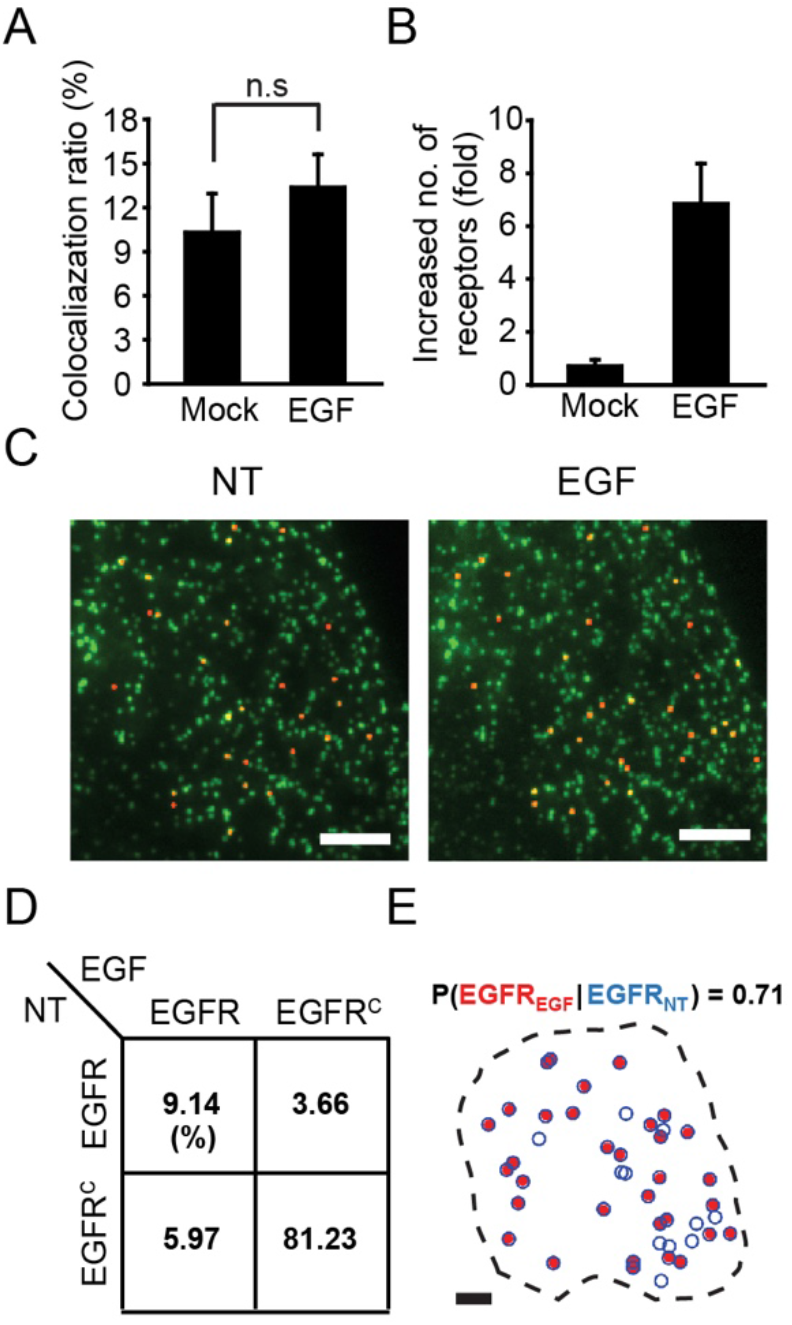
Related to Figure 5. Pre-allocated subset of CCP for EGFR primarily utilized after EGF stimulation. (A) Diffusion-based molecular colocalization analysis of CCPs with EGFR with EGF treatment in a single live BSC-1 cell. (B) The quadrants of CCP ratios that contain EGFR or do not contain EGFR before and after EGF treatment. (C) Maps showing conditional probability of the CCP subsets containing EGFR after mock treatment, given the subset before EGF treatments. Black dashed lines display cell boundaries, blue circles indicate the CCP positions before the respective treatments, and red dots indicate the CCP positions after the respective treatments. (D, E) Histograms show the colocalization fraction of CCP with EGFR (D) and fold increase of the number of EGFRs colocalized with CCPs (F) after mock or EGF treatment. Error bars denote s.e.m. at the single-cell level (n > 5).

**Figure S7.**
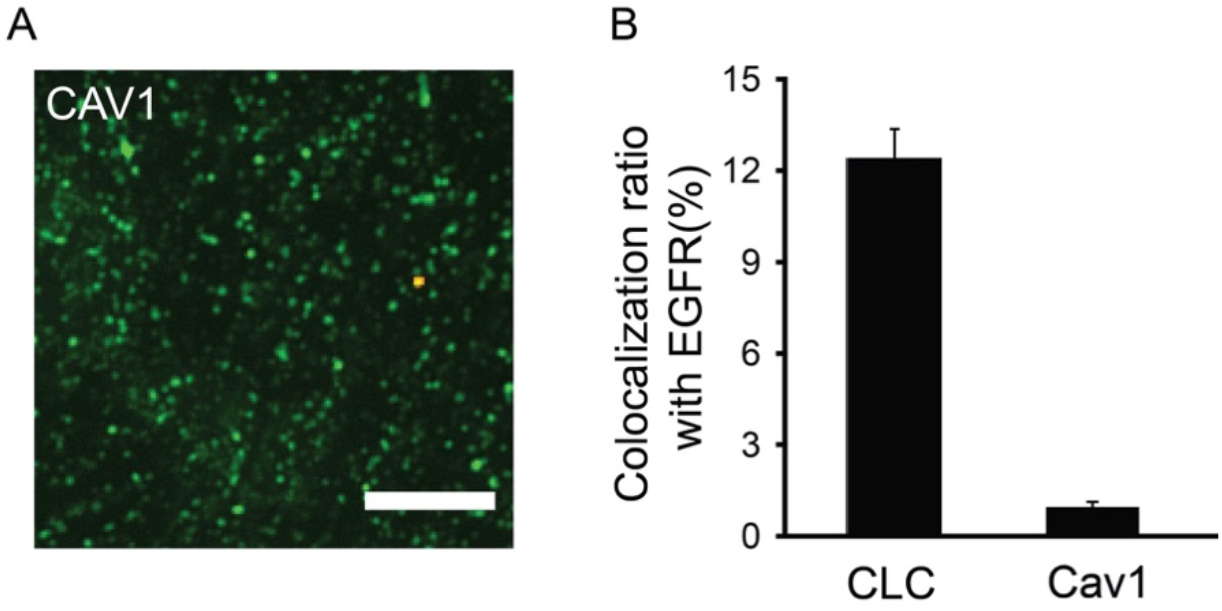
Related to Discussion. Cav1 colocalization with EGFR in live mEGFP-Cav1 expressing BSC1 cell line. (A) Diffusion-based single-molecule colocalization image of mEGFP-Cav1 and anti-EGFR Fab fragment labeled with Alexa 647. Green and red dots represent Cav1 without EGFR and Cav1 with EGFR, respectively. Scale bar, 5μm. (B) A histogram shows the colocalization ratio of indicated molecular complex with EGFR. Error bars for denote s.e.m. at the single-cell level (n > 3).

**Movie S1.**
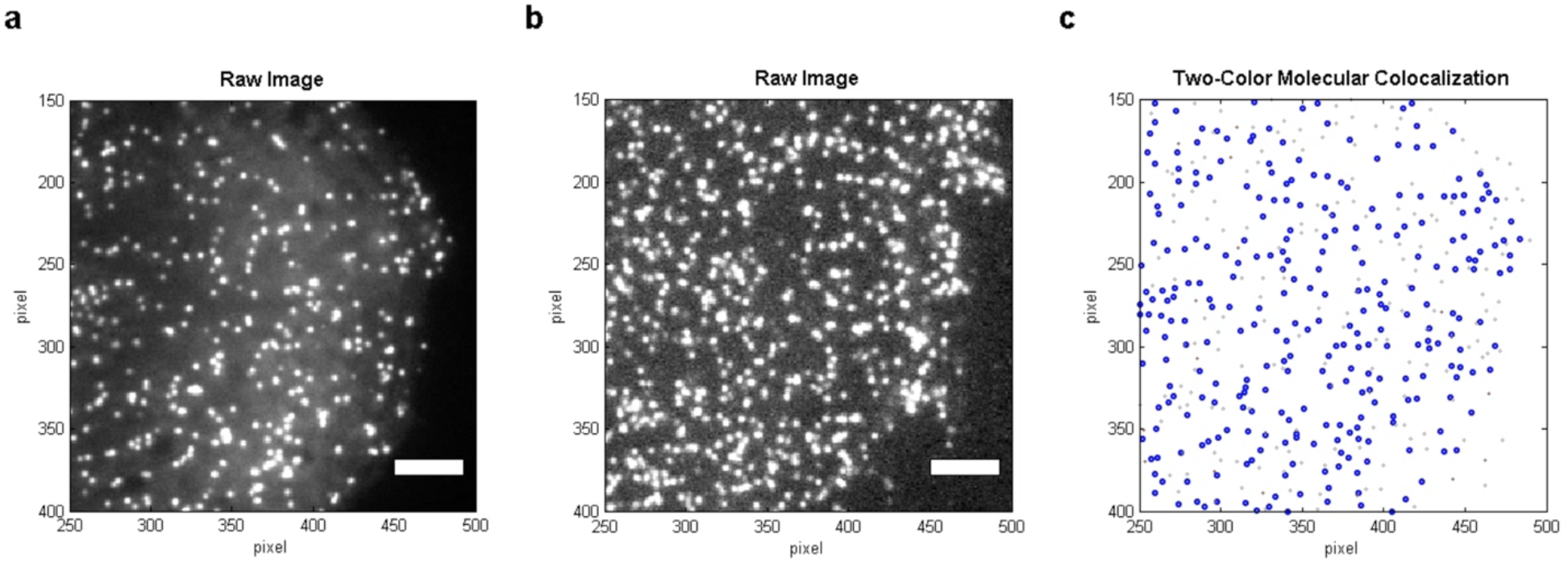
Related to Figure 1. Diffusion-based molecular colocalization between CCSs and single-molecule EGFRs. (A) The raw movie of EGFP-CLC acquired using TIRFM at 0.5 Hz. (B) The raw movie of single-molecule EGFR visualized using dSTORM with an Alexa Fluor 647-labeled Fab fragment of an anti-EGFR antibody acquired at 20 Hz acquired in a single live BSC-1 cell. (C) Diffusion-based molecular colocalization analysis. Blue circles indicate CCSs, black lines indicate single-molecule EGFR trajectories, and red dots indicates single-molecule EGFR trajectories matching to the diffusivity of CCPs. Scale bars, 5 μm.

**Movie S2.**
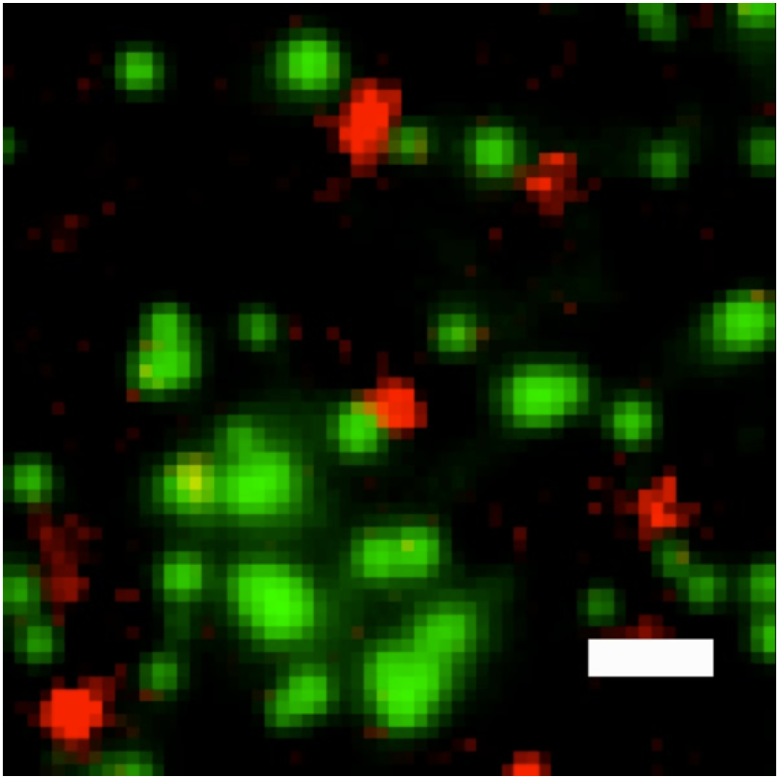
Related to Figure 3. Transient trapping of single-molecule EGFR into a CCP in a resting state. The overlay movie of EGFP-CLC and single-molecule EGFR visualized using dSTORM with an Alexa Fluor 647-labeled Fab fragment of an anti-EGFR antibody acquired at 20 Hz in a single live BSC-1 cell. Two-color channels were registered with a > 10-nm resolution. Scale bars, 1 μm.

**Movie S3.**
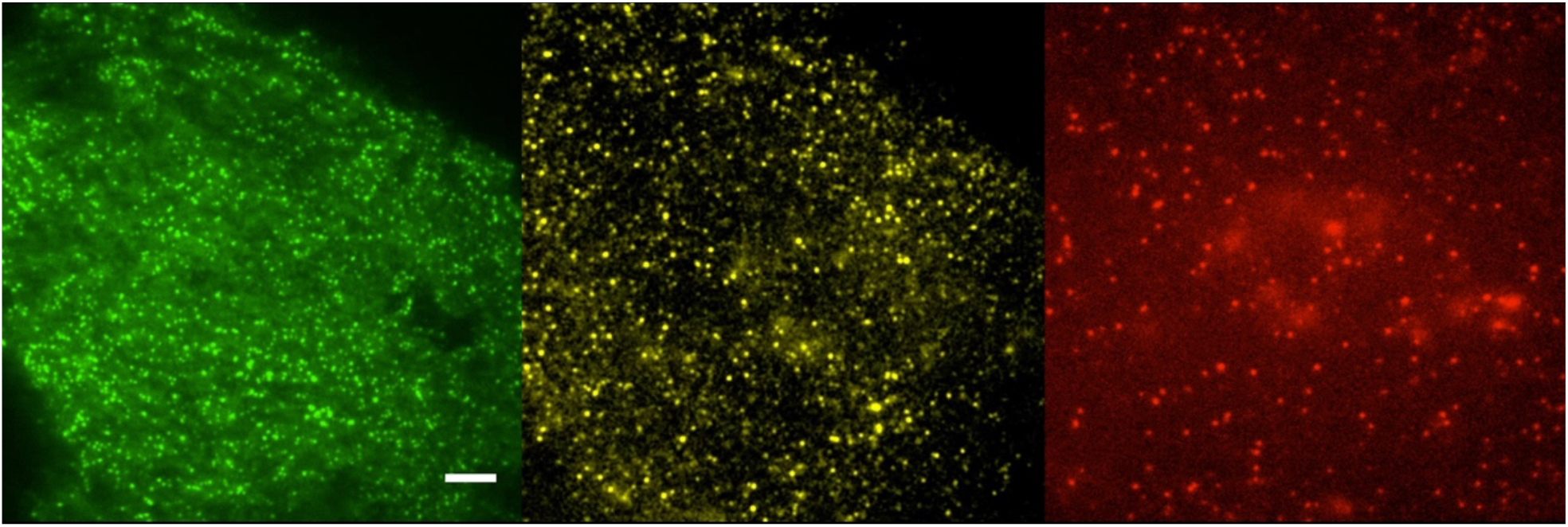
Related to Figure 5. Three-color molecular colocalization among CCPs, single-molecule TfR, and single-molecule EGFR. The raw movie of EGFP-CLC, Cy3b-labeled SNAP-TfR and Alexa Fluor 647-labeled endogenous EGFR acquired using TIRFM at 20 Hz in a single live BSC-1 cell. Three-color channels were registered with a > 15-nm resolution. Scale bars, 5 μm.

